# Reduced S-nitrosylation of TGFβ1 elevates its binding affinity towards the receptor and promotes fibrogenic signaling in the breast

**DOI:** 10.1101/2023.09.07.556714

**Authors:** Joshua Letson, Saori Furuta

## Abstract

Transforming Growth Factor β (TGFβ) is a pleiotropic cytokine closely linked to tumors. TGFβ is often elevated in precancerous breast lesions in association with epithelial-to-mesenchymal transition (EMT), indicating its contribution to precancerous progression. We previously reported that basal nitric oxide (NO) levels declined along with breast cancer progression. We then pharmacologically inhibited NO production in healthy mammary glands of wild-type mice and found that this induced precancerous progression accompanied by desmoplasia and upregulation of TGFβ activity. In the present study, we tested our hypothesis that NO directly S-nitrosylates (forms an NO-adduct at a cysteine residue) TGFβ to inhibit the activity, whereas the reduction of NO denitrosylates TGFβ and de-represses the activity. We introduced mutations to three C-terminal cysteines of TGFβ1 which were predicted to be S-nitrosylated. We found that these mutations indeed impaired S-nitrosylation of TGFβ1 and shifted the binding affinity towards the receptor from the latent complex. Furthermore, *in silico* structural analyses predicted that these S-nitrosylation-defective mutations strengthen the dimerization of mature protein, whereas S-nitrosylation-mimetic mutations weaken the dimerization. Such differences in dimerization dynamics of TGFβ1 by denitrosylation/S-nitrosylation likely account for the shift of the binding affinities towards the receptor vs. latent complex. Our findings, for the first time, unravel a novel mode of TGFβ regulation based on S-nitrosylation or denitrosylation of the protein.

**Significance statement:** Transforming Growth Factor β (TGFβ) is a widely studied cytokine associated with tumors. Because of its pleiotropic functions and dichotomous roles in tumorigenesis, the development of therapeutics targeted to TGFβ for cancer treatment has been challenging. In the present study, we report that TGFβ is indeed S-nitrosylated at specific sites for repressing its functions, whereas it is denitrosylated to derepress its activity. Such covalent modification-based regulation of TGFβ activity could potentially be utilized to design a new type of inhibitor or activator of the protein.

## Introduction

Formation of dense collagenous stroma, termed desmoplasia, is often detected as a hard ‘lump’ and is a hallmark of cancer (1). Desmoplasia, however, may actually happen in non- or pre-cancerous breast tissues, serving as a strong risk factor for breast cancer (2). Desmoplasia is attributed to the emergence of myofibroblasts (MyoFbs) which are highly contractile and secretory stromal cells derived from various types of tissue resident cells. They are present in cancerous as well as precancerous and tumor-adjacent normal tissues and proposed as a key player in tumor initiation (3, 4). MyoFb differentiation is primarily induced by paracrine factors secreted from parenchymal and stromal cells, namely, Transforming Growth Factor β (TGFβ) (5). TGFβ is a family of cytokines that exert pleiotropic functions (6). For its relevance to tumors, however, TGFβ plays complex and dual roles, serving as both a tumor-suppressor and -promoter depending on the stage and tissue type of tumorigenesis (7). TGFβ expression is often elevated in precancerous breast lesions in association with epithelial-to-mesenchymal transition (EMT) (8, 9), suggesting its possible role in the formation of MyoFbs in these lesions (10).

To determine the mechanism of precancerous progression of the breast, we previously inhibited nitric oxide (NO) in healthy mammary glands of wild-type mice (11, 12). NO is a bioactive signaling molecule produced throughout the body, and its aberrant production is linked to different risk factors of breast cancer (13–26). We found that the basal levels of NO plummeted along with breast cancer progression. Importantly, we saw that pharmacological deprivation of NO in healthy mammary glands induced precancerous progression accompanied by desmoplasia and TGFβ upregulation (11, 12). These findings suggested that NO plays critical roles in suppressing TGFβ activity and that its deficit could lead to precancerous progression of the breast. In the present study, we explored the mechanism by which TGFβ activity is suppressed by NO. We hypothesized that TGFβ is directly targeted for S-nitrosylation (SNO), an NO-mediated covalent modification of cysteine residues, which would induce certain conformational changes and modulate protein functions (27, 28). To test this possibility, we introduced mutations to three C-terminal cysteines of TGFβ1 which were predicted to be S-nitrosylated. We found that mutations at these sites indeed impaired SNO of TGFβ1 and shifted their binding affinity towards the receptor from the latent complex. Further conformational analyses predicted that these SNO-defective mutations would induce stronger dimerization of mature TGFβ1, whereas SNO at these sites would weaken dimerization, possibly accounting for the shifts of the binding affinities. Our results unravel a novel mechanism of regulating TGFβ activity through SNO or denitrosylation that impacts dimerization and protein-protein interactions.

## Results

### NO downregulates TGFβ activity in normal mammary epithelial cells

We previously reported that pharmacological inhibition of basal nitric oxide (NO) production in wild-type mice induced precancerous progression of mammary glands accompanied by highly desmoplastic stroma. These glands also showed elevated TGFβ activity and fibrogenic signaling (11, 12). To explore the mechanism of this phenomenon, we modulated NO levels *in vitro* using cell lines of MCF10A human breast cancer progression series comprising normal MCF10A and malignant CA1d cells (29). To this end, we applied a NOS inhibitor, L-NAME (2.5 mM), NOS agonist, L-arginine (L-ARG, 2.5 mM), or NO donor SNAP (10 µM) to cultured MCF10A cells and determined the levels of TGFβ1 and its downstream effector, SMAD3 (phosphorylated and total SMAD3). We saw a strong negative correlation of NO levels to TGFβ levels and SMAD3 activation. L-NAME greatly elevated both the full-length and mature form of TGFβ as well as SMAD3 activation (pSMAD3), whereas both L-ARG and SNAP downmodulated all of them (**Fig. 1A-C**). Consistently, phosphorylation of the receptor, TGFβR1, was concordant with phosphorylation of SMAD3. The same impacts of NO modulators on the receptor activation were observed in both MCF10A and CA1d cells (**Fig. 1D**).

**Figure 1:**
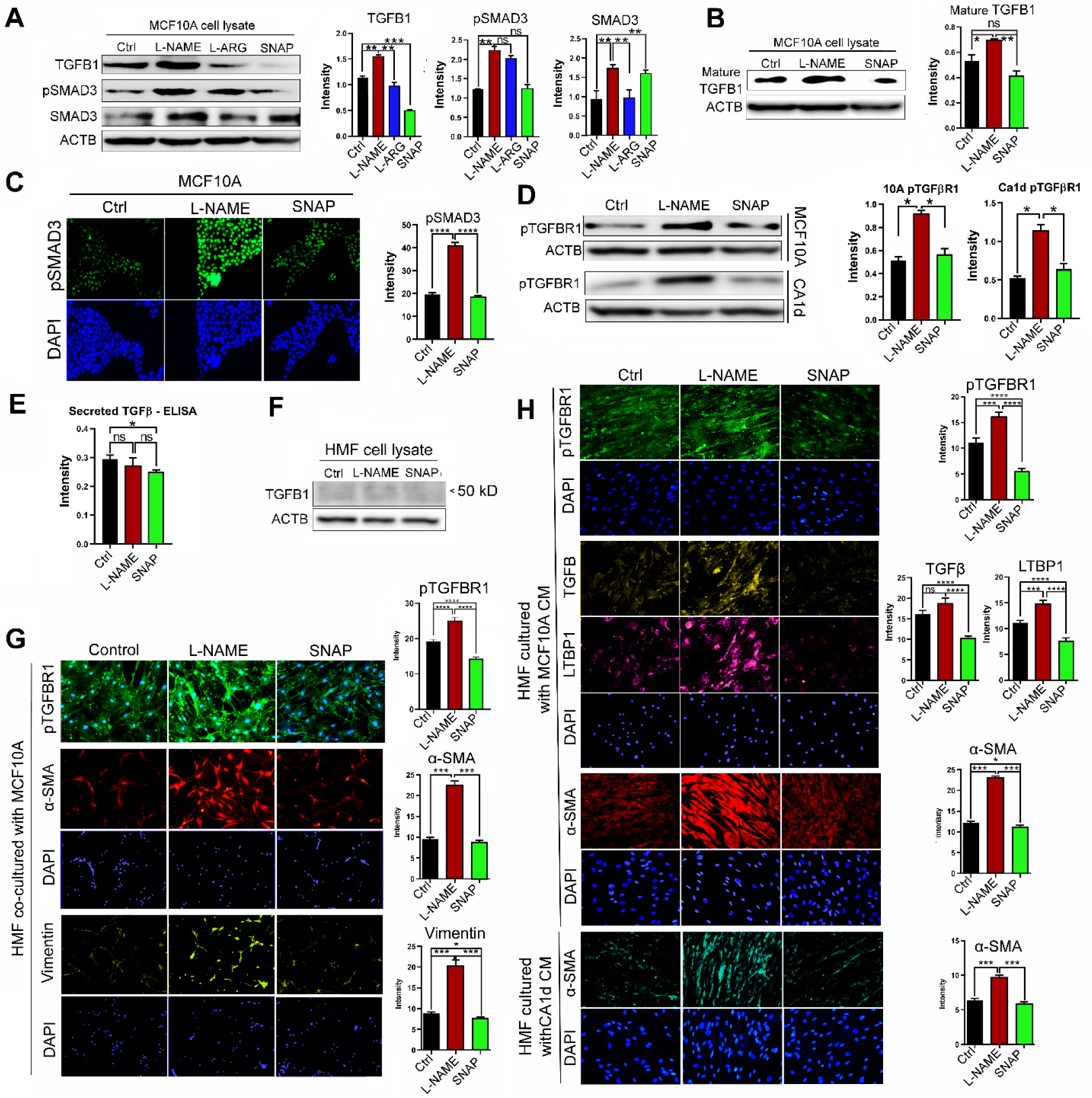
Basal NO production in mammary epithelial cells suppresses TGFβ1 signaling. **A**) (Left) Western blot (WB) showing the protein expression levels of the full-length TGFβ1 and its downstream effector, pSMAD3, in normal human mammary epithelial cells (MCF10A) following treatment with NOS inhibitor, L-NAME (2.5 mM); NOS agonist, L-arginine (2.5 mM); or NO donor, SNAP (10 μM) for 24 hours. β-actin (ACTB) was used as an internal control. (Right) Quantification of TGFβ1, pSMAD3 and SMAD3 levels. **B**) (Left) Western blot showing the levels of mature TGFβ1 in MCF10A cells after treatment with L-NAME or SNAP. (Right) Quantification of mature TGFβ1 levels. **C**) (Left) MCF10A cells stained with anti-pSMAD3 antibody after treatment with L-NAME or SNAP. (Right) Quantification of pSMAD3 signals. **D**) (Left) WB showing levels of activated TGFβ1 receptor, pTGFBR1, in MCF10A cells (top) and isogenic cancer cells CA1d (bottom) after treatment with L-NAME or SNAP. (Right) Quantification of pTGFBR1 signals in both cell lines. **E**) ELISA detection of the level of mature TGF β1 secreted from MCF10A cells after L-NAME or SNAP treatment. **F**) Measurement of TGFβ1 levels in HMFs following treatment with L-NAME or SNAP. Note that there is no expression of TGFβ1 in HMFs alone even after treatment with NO modulators. **G**) (Left) HMFs co-cultured with MCF10A cells in the presence of L-NAME or SNAP and stained with antibodies against molecules downstream of TGFβ1 signaling. (Right) Quantification of TGFβ1 signaling molecules. **H**) HMFs cultured with the conditioned media (CM) of MCF10A cells treated with L-NAME or SNAP and stained with antibodies against molecules downstream of TGFβ1 signaling. (Right) Quantification of TGFβ1 signaling molecules. Error bars: ± SEM. *, p ≤ 0.05; **, p ≤ 0.01; ***, and p ≤ 0.001; p >0.05 was considered significant.

Next, we sought to determine the paracrine signaling of TGFβ in response to modulation of NO levels. The media of drug-treated MCF10A cells showed a significant decrease of secreted TGFβ level after SNAP treatment (**Fig. 1E**). To test the functionality of secreted TGFβ, we co-cultured the drug-treated MCF10A cells with human mammary fibroblasts (HMFs). HMFs alone did not express TGFβ and did not respond to NO modulators (**Fig. 1F**). However, HMFs co-cultured with L-NAME-treated MCF10A cells showed significant increases of phospho-TGFβR1 and downstream fibrogenic proteins, α-smooth muscle actin (αSMA) and vimentin. Conversely, HMFs co-cultured with SNAP-treated MCF10A cells showed significant decrease in these proteins (**Fig. 1G**). Alternatively, we cultured HMFs with the conditioned media (CM) from the drug-treated MCF10A cells. Consistent with the co-culture system, CM of L-NAME-treated MCF10A cells largely elevated TGFβ signaling molecules, whereas CM of SNAP-treated MCF10A cells downmodulated these proteins (**Fig. 1H**).

### TGFβ1 is S-nitrosylated at specific cysteines to suppress the activity, whereas SNO-defective mutants show increased activity

To explore the mechanism by which NO suppresses TGFβ activity, we tested our hypothesis that TGFβ is directly targeted for S-nitrosylation (SNO), an NO-mediated covalent modification of cysteines, to suppress its functions. SNO takes place in over 3000 proteins that meet specific physical and biochemical requirements, inducing conformational changes to modulate the protein functions (27, 30). Potential candidate proteins and specific SNO sites could be predicted using software such as GPS-SNO (31). Upon inputting the full-length TGFβ1 protein (390 AAs) sequence into GPS-SNO, the program predicted three C-terminal cysteines, C355, C356, and C389, as the potential SNO sites with high probabilities (**Fig. 2A, B**). To validate whether these sites are indeed S-nitrosylated, we introduced point mutations by substituting cysteines with alanines (C355A, C356A, and C389A) and expressed these mutants in comparison to wild-type TGFβ1 and an empty vector (PCDF1) control in MCF10A cells. Overexpression of these ectopic TGFβ1 proteins were confirmed by RT-PCR (**Fig. 2C**). We also used siRNAs that specifically targeted the endogenous TGFβ1 to confirm the strong expression of the ectopic proteins (**Table S1**, **Fig. S1A, B**). Then, we determined S-nitrosylation levels of these overexpressing cell lines using iodoTMT-tagging/pull-down method (32). We indeed saw strong SNO of both full-length and mature TGFβ1 in cells overexpressing the wild-type protein, whereas SNO levels of TGFβ1 significantly declined in cells overexpressing three mutant constructs (**Fig. 2D**). These results strongly indicate that TGFβ1 is S-nitrosylated at these three cysteines.

**Figure 2:**
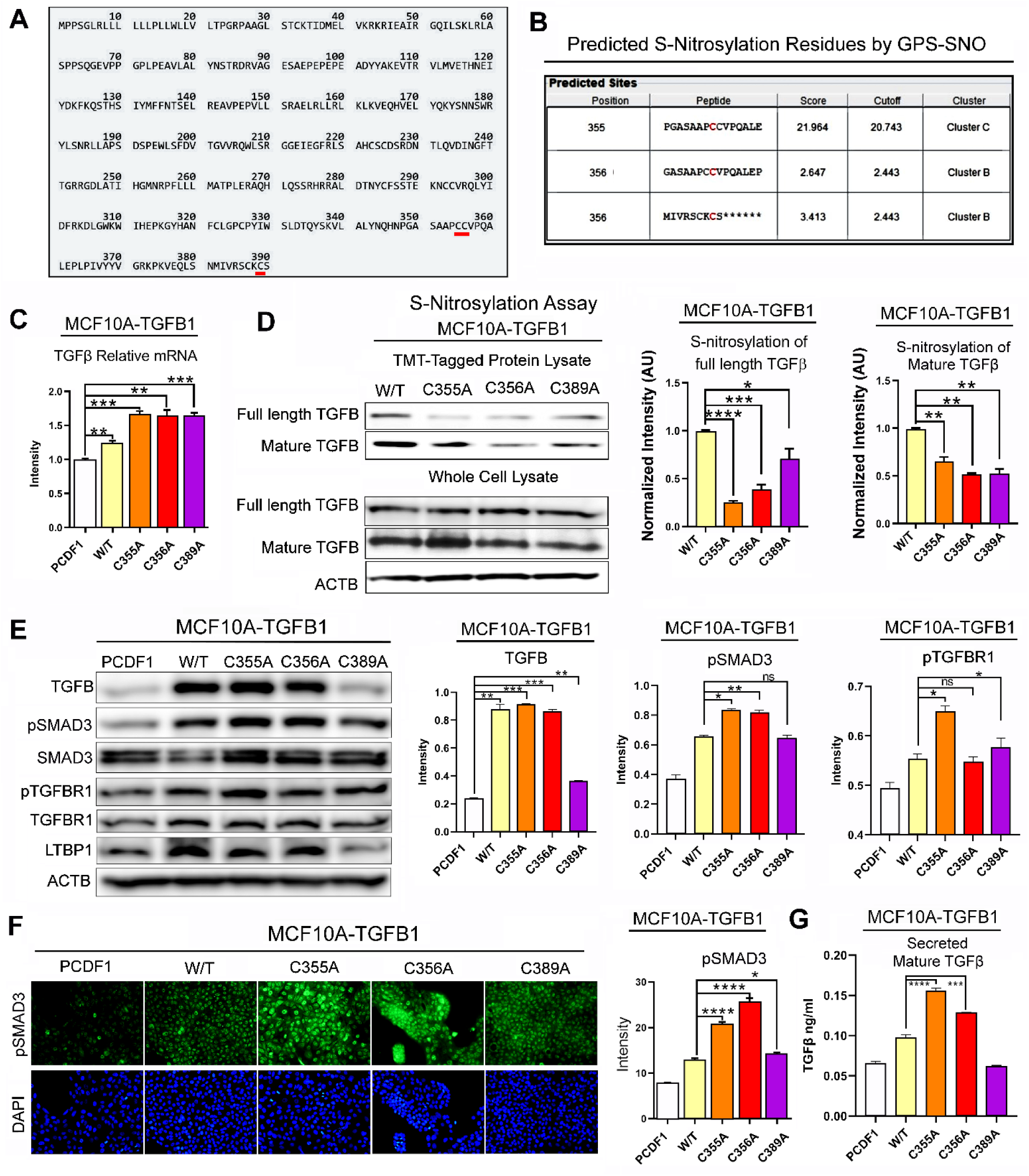
Impaired S-nitrosylation of TGFβ1 activates its downstream signaling. **A**) Full-length TGFβ1 protein sequence showing three predicted S-nitrosylation (SNO) sites (highlighted). **B**) Probability scores of three cysteines predicted to be SNO sites by GSP-SNO software. **C**) TGFβ1 transcripts in MCF10A cells overexpressing wild-type (WT) or SNO-defective mutants (C355A, C356A, or C389A) of TGFβ1. Note that TGFβ1 transcript levels increased in all the overexpressing cell lines, confirming the expression of the ectopic TGFβ1. **D**) (Left) Determination of SNO levels in TGFβ1 overexpressing MCF10A cell lines (W/T or SNO-defective mutants) by iodoTMT-labeling/pull-down and western blotting against TGFβ1 (full-length and mature). (Right) Quantification of SNO levels of full-length and mature TGFβ1. NOTE that mutant TGFβ1-expressing cell lines showed significantly reduced SNO levels of both full-length and mature TGFβ1. **E**) (Left) Western blots showing the levels of TGFβ1 signaling molecules in TGFβ1-overexpressing MCF10A cell lines (WT or mutants). ACTB was used as an internal control. (Right) Quantification of the levels of TGFβ1 signaling molecules in TGFβ1-overexpressing cell lines. **F**) (Left) TGFβ1-overexpressing cell lines (WT or mutants) stained with an antibody against pSMAD3. (Right) Quantification of pSMAD3 levels in TGFβ1-overexpressing cell lines. **G)** ELISA detection of mature TGFβ1 secreted from MCF10A W/T or SNO-defective mutants. Error bars: ± SEM. *, p ≤ 0.05; **, p ≤ 0.01; ***, and p ≤ 0.001; p >0.05 was considered significant.

Next, we sought to investigate the effects of SNO vs. denitrosylation of these sites on downstream signaling. We saw significant increases in TGFβ1 levels and downstream signaling molecules (pSMAD3 and pTGFBR1) in cells overexpressing SNO-defective (denitrosylated) TGFβ1 mutants (**Fig 2 E, F**). Secreted levels of TGFβ were also significantly elevated in SNO-defective cell lines compared to the wild-type construct (**Fig. 2G**). Consistently, fibrogenic molecules, αSMA and vimentin, were also elevated in MCF10A cells overexpressing either of the three mutants (**Fig. 3A**) as well as HMFs cultured with CM of these MCF10A cell lines (**Fig. 3B**). To verify that such induction of fibrogenic signaling was indeed caused by TGFβ1, these MCF10A cell lines were treated with a TGFβ inhibitor, galunisertib (Gal). As expected, the induction of fibrogenic signaling was compromised by Gal treatment, further attesting to the critical roles of SNO-defective TGFβ1 in fibrogenic activation (**Fig. 3C**).

**Figure 3:**
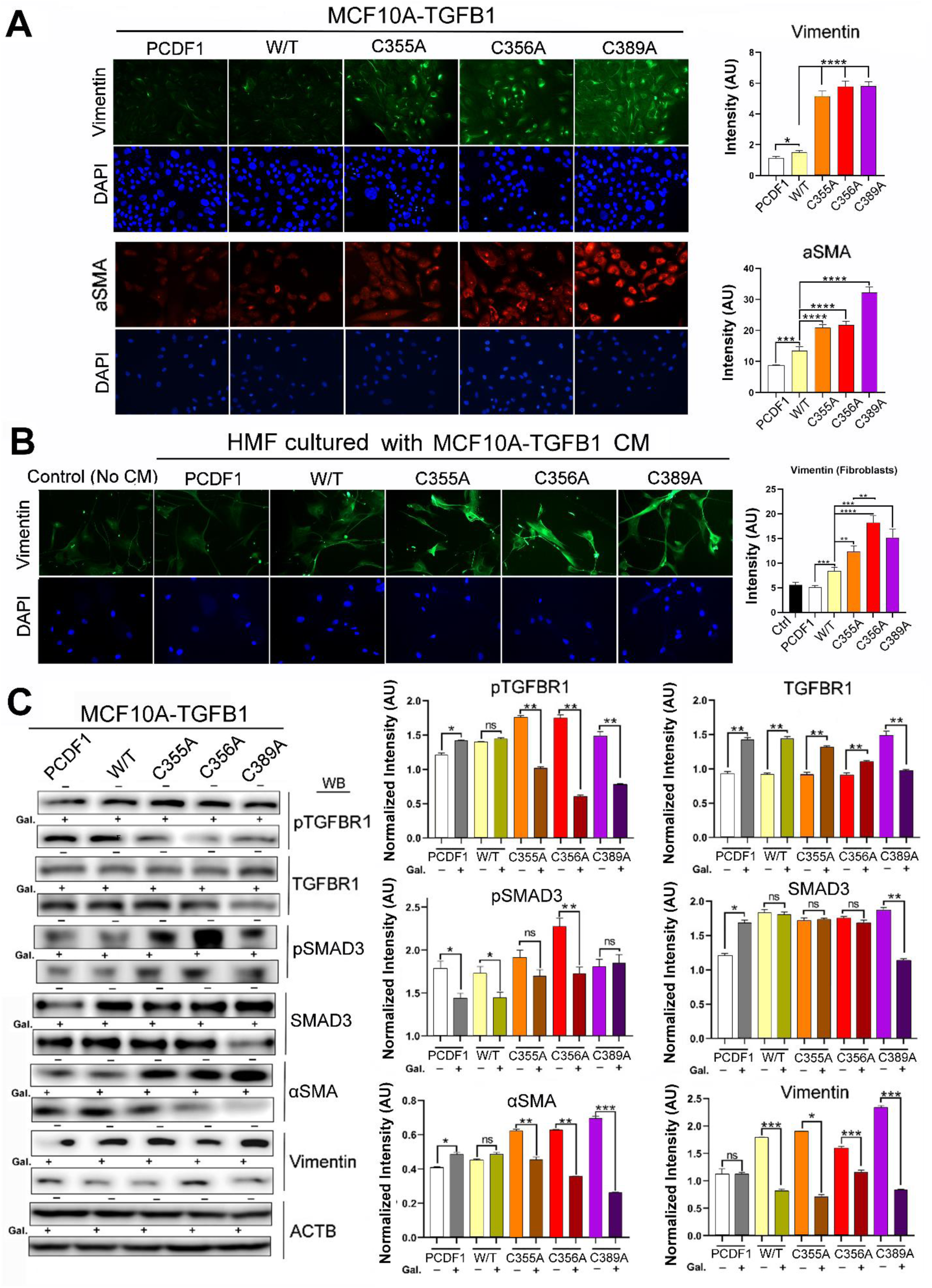
Impaired S-nitrosylation of TGFβ1 promotes fibrogenic signaling. **A**) (Left) TGFβ1-overexpressing MCF10A cell lines (WT or SNO-defective mutants) stained with antibodies against fibrogenic markers, vimentin and α-smooth muscle actin (αSMA). (Right) Quantification of vimentin and αSMA levels in TGFβ1-overexpressing cell lines. Note the significant increases in both fibrogenic markers in SNO-defective mutants. **B**) (Left) HMFs cultured with the CM of TGFβ1-overexpressing MCF10A cell lines (WT or mutants) and stained with an antibody against vimentin. (Right) Quantification of vimentin levels in TGFβ1-overexpressing cell lines. **C**) (Left) Western blot for the validation of the impacts of TGFβ1-overexpression on fibrogenic signaling using treatment with a TGFβ1 inhibitor galunisertib (Gal, 10 µM, 48 hours). ACTB was used as an internal control. (Right) The levels of fibrogenic molecules normalized against Gal-untreated samples in TGFβ1-overexpressing cell lines with or without Gal treatment. Error bars: ± SEM. *, p ≤ 0.05; **, p ≤ 0.01; ***, and p ≤ 0.001; p >0.05 was considered significant.

### SNO of TGFβ1 shifts its binding affinities towards the latent complex, while SNO-defective mutants have affinities towards the receptor due to the difference in dimerization dynamics

To explore the mechanistic basis for SNO-mediated suppression of TGFβ1 activities, we analyzed the interactions of the wild-type vs. SNO-defective TGFβ1 mutants with the latent complex protein, LTBP1, and the receptor, TGFBR1. We saw that SNO-defective mutants showed significantly reduced binding with LTBP1, but increased binding with TGFBR1 (both the total and phosphorylated proteins) (**Fig. 4A**). This result suggests that SNO of TGFβ1 shifts its binding affinity towards the latent complex, whereas denitrosylation (SNO-defective mutations) of TGFβ1 shifts the affinity towards the receptor.

**Figure 4:**
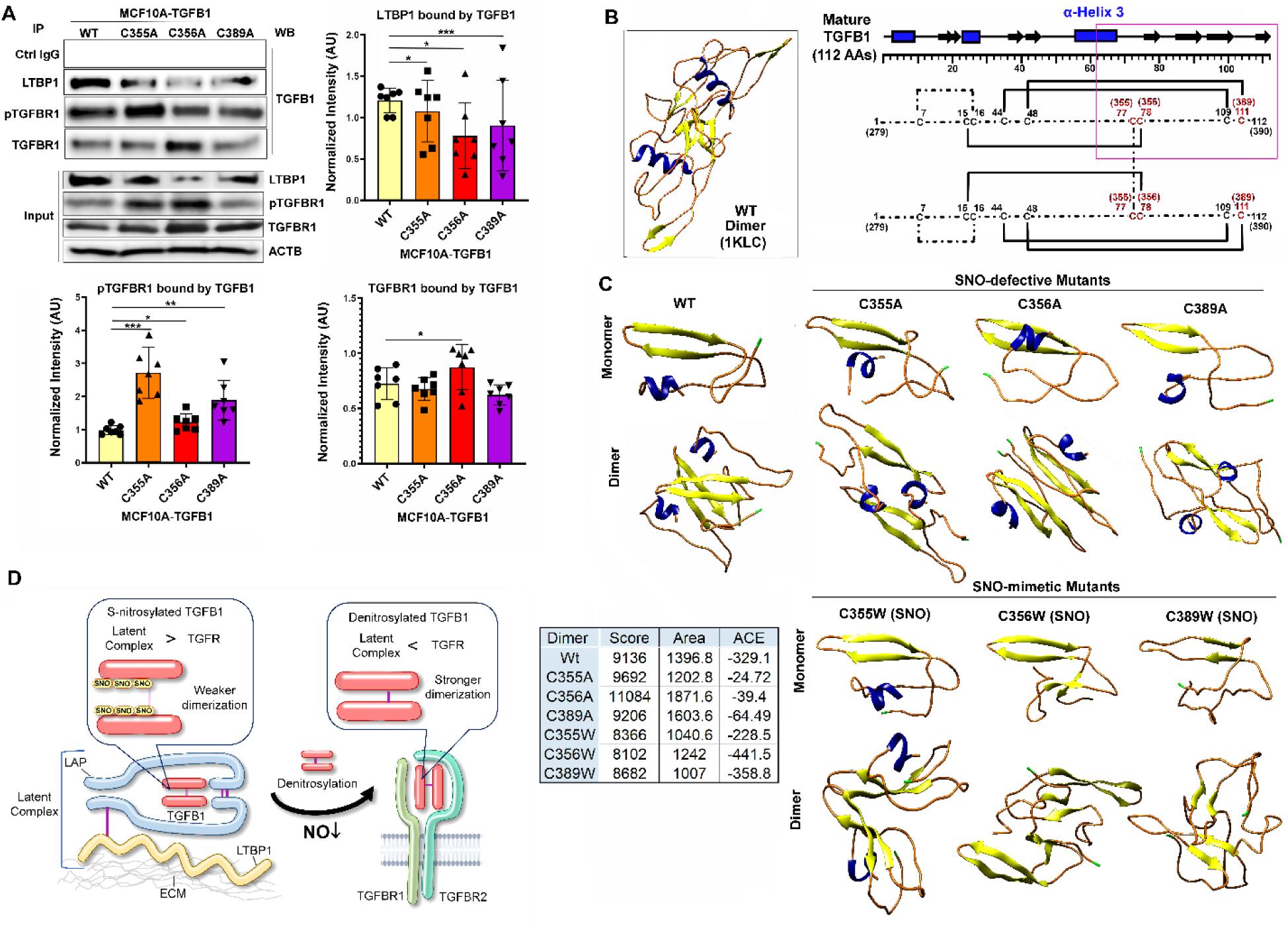
Impaired S-nitrosylation of TGFβ1 elevates its binding affinities towards the receptor over the latent complex through a change in dimerization dynamics. **A**) (Left top) Immuno-precipitation to determine the interaction of W/T or mutant TGFβ1 with the latent complex (LTBP1) vs. receptor (pTGFBR1 and TGFBR1). (Left middle) Western blot showing the input levels of LTBP1, pTGFBR1, and TGFBR1. ACTB was used as an internal control. (Right) The levels of LTBP1, pTGFBR1 and TGFBR1 normalized against the input levels in TGFβ1-overexpressing cell lines. Error bars: ± SEM. *, p ≤ 0.05; **, p ≤ 0.01; ***, and p ≤ 0.001; p >0.05 was considered significant. **B**) (Left) The structure of mature TGFβ1 dimer (PDF ID: 1KLC) featuring α-helix 3 (blue) and β-strands (yellow). Note the symmetrical conformation and aligned α-helices. (Right) Schematic representation of the dimer of mature TGFβ1 (112AAs) featuring intra- and inter-chain disulfide linkages. AA numbers are shown as both the numbers in mature and full-length proteins (parenthesized). SNO sites are shown in red. Boxed region: C-terminal 50 AAs were analyzed for fold prediction (PEP-FOLD3) and dimerization (SymmDoc). **C**) (Top) Predicted monomeric or dimeric structures of the C-terminal 50 AAs of mature WT and SNO-defective mutants (C355A, C356A, or C389A) of TGFβ1. Note the symmetrical conformations and aligned α-helices of all the proteins. (Bottom) Predicted monomeric or dimeric structures of the C-terminal 50 AAs of mature WT and SNO-mimetic mutants (C355W, C356W, or C389W) of TGFβ1. Note the distanced α-helices of C355W mutant dimer and loss of α-helices in C356W and C389W mutant dimers. (Table) Dimerization parameters of WT and mutant TGFβ1 proteins. Score: Shape complementarity score (36); Area: Interface area; and ACE: Atomic contact energy (37). **D**) Working model of the SNO-mediated affinity shift of mature TGFβ1, where SNO shifts the affinity toward the latent complex and denitrosylation shifts it towards the receptor.

To further investigate the mechanistic basis of this phenomenon, we sought to compare the three-dimensional (3D) conformations of the wild-type vs. SNO-defective TGFβ1 mutants. Wild-type mature TGFβ1 (112 AAs, PDB ID: 1KLC) formed a symmetrical dimer characterized by centrally clustered β-strands (β-sheets) and parallelly aligned α-helices on the side (**Fig. 4B, left**). In particular, interactions between two α-helices 3, located in the middle of mature TGFβ1 (**Fig. 4B, right**). play essential roles in the enhanced rigidity and stability of the dimer. Stable dimerization would be critical for tight binding to the receptor for the induction of downstream signaling (33).

To determine the contributions by the three cysteines targeted for SNO, we generated the *in silico* 3D conformation of the 50 AA region between α-helix 3 and the C-terminus of mature TGFβ1 (**Fig. 4B, right**) using PEP-FOLD3 software (34). Then, a monomer with the highest probability score was assembled into a homodimer using SymmDoc software based on the most favorable dimerization parameters (35). To validate the feasibility of this approach, the wild-type sequence was assembled into a homodimer and compared to the published structure (1KLC) (**Fig. 4B, right**). As expected, the wild-type sequence was predicted to form a dimer that resembled the published structure featuring axial symmetry with centrally stacked β-sheets and flanking parallel α-helices (**Fig. 4C, top)**. All three SNO-defective mutants (C355A, C356A and C389A), on the other hand, were predicted to form more compact dimers that retained the axial symmetries and closely aligned α-helices, but had two β-sheets more distantly localized than the wild-type. Because of the proximity of two α-helices in these mutants, their dimerization, and thus, interactions with the receptor would be stronger than the wild-type (33). Stronger dimerization of the three SNO-defective mutants was in fact reflected by their increased values of shape complementarity scores and atomic contact energies (ACE) indicative of hydrophobicity (**Fig. 4C, table**)(36, 37). We next analyzed SNO-mimetic mutants where the three reactive cysteines were changed to tryptophans (C355W, C356W, and C389W) (38, 39). C355W mutant was predicted to form a more spread-out dimer, although it retained the axial symmetry with two parallel α-helices and central β-sheets. These secondary structures in one monomer were distanced from those in the other monomer, indicating looser dimerization. More dramatically, C355W and C389W mutants were predicted to form disorganized dimers that lost parallel α-helices (**Fig. 4C, bottom**). Because of the lack of α-helices, their dimerization and receptor binding would be weaker than the wild-type (33). Weaker dimerization of the three SNO-mimetic mutants was reflected by their reduced values of shape complementarity scores and ACE (**Fig. 4C, table**)(36, 37). These shifts in dimerization dynamics through SNO (SNO-mimetic mutations) vs. denitrosylation (SNO-defective mutations) may account for the differences in the binding affinities of mature TGFβ1 towards the latent complex vs. receptor (**Fig. 4A**).

## Discussion

It has been reported that mature TGFβ proteins alter their conformations between ‘open’ and ‘closed’ forms in solution. The closed, compact conformation is formed once two α-helices 3 in two monomers establish stable interactions and is a favorable condition for receptor binding. However, the predominance of either state depends on a specific TGFβ isomer. For example, TGFβ1 prefers closed over open conformations, whereas TGFβ3 prefers open over closed conformations (33). Interestingly, these biochemical differences in the closed vs. open conformations of TGFβ1 are analogous to the differences we saw in the structures and functionalities of denitrosylated (SNO-defective mutant) vs. S-nitrosylated (SNO-mimetic mutant) TGFβ1. It is thus possible that SNO of mature TGFβ1 helps shift the conformational equilibrium towards the otherwise unfavorable open conformation to weaken the binding affinity towards the receptor. This speculation is in part supported by a report that many cysteines (including those targeted for SNO) of TGFβ1 are critical for the latency and their loss could lead to the constitutive activation of the protein (40).

Our previous and present studies strongly suggest that TGFβ1 is highly S-nitrosylated for latency in normal breast cells. However, as the basal NO production declines possibly due to the increase in oxidative stress, TGFβ1 becomes denitrosylated and derepressed, contributing to the induction of desmoplasia and precancerous progression (11, 12, 27). Roles of TGFβ1 in precancerous progression, however, remain controversial. Classically, TGFβ1 is proposed to exert tumor suppressive roles in the early-stage carcinogenesis. In contrast, some more recent studies demonstrated that TGFβ1 acts to stabilize precancerous cells by preventing p53-mediated apoptosis and other stress responses (7–9, 41–43). Our studies also strongly indicate that de-repression of TGFβ1, in response to the impairment of basal NO production in the breast, has positive roles in precancerous progression through induction of fibrogenic signaling. Our observation is in line with well-established roles of TGFβ in EMT-mediated tumorigenesis (8, 9).

Reduced NO bioavailability is associated with different chronic conditions, such as diabetes, cardiovascular disease, and obesity, as well as aging. It often leads to the formation of stiff vasculature and desmoplastic extracellular matrix (ECM), which is directly linked to increased cancer risk (44–46). Complementarily, moderate to leisure time exercise is shown to elevate basal NO production, ameliorate vascular stiffening, and reduce cancer risks of older people (47–49). Such NO-associated tissue stiffening is largely attributed to the downmodulation of SNO-mediated inhibition of ECM-remodeling enzymes, including tissue transglutaminases and matrix metalloproteinases (27, 47). In addition, our study reports that a major fibrogenic protein, TGFβ1, is another contributor to tissue stiffening induced by reduction of SNO (11, 12). While in healthy cells a subset of proteins are constitutively S-nitrosylated to remain under control, in disease states, on the other hand, SNO levels could be dysregulated, contributing to disruption of homeostasis (30). Development of novel therapeutics aimed to modulate SNO levels of specific proteins may warrant further investiations.

## Materials and Methods

### Cell Lines

Throughout the study, the MCF10A breast cancer progression series cell lines were utilized consisting of MCF10A and Ca1d and were obtained from Karmanos Cancer Institute (MI, USA). Primary human mammary fibroblasts (HMFs) were obtained from ScienCell (ScienCell Research Laboratories, Carlsbad, CA, USA, Cat. # 7630). 293FT cells were obtained from Thermo Fisher Scientific (Waltham, MA, USA Cat. # R70007).

### Cell Culture

The MCF10A and Ca1d cell lines were cultured in DMEMF12 (Thermo Fisher Scientific, Waltham, MA, USA, Cat. # 11320033) and supplemented with the following: 5% horse serum (Thermo Fisher Scientific, Waltham, MA, USA, Cat. # 16050122); 1% penicillin/streptomycin (Thermo Fisher Scientific, Waltham, mA, USA, Cat. # 15140122); 20 ng/mL EGF (Sigma, St. Louis, MO, USA, Cat. # E-9644); 10 μg/mL insulin (Sigma, St. Louis, MO, USA, Cat. # I-1882); 100 ng/mL cholera toxin (Sigma, St. Louis, MO, USA, Cat. # C-8052) and 0.5 μg/mL hydrocortisone (Sigma, St. Louis, MO, USA, Cat. # H-0888). The HMF cell line was cultured in fibroblast basal medium (Sigma, St. Louis, MO, USA, Cat. #115-500) and supplemented with 1% fibroblast growth supplement (Sigma, St. Louis, MO, USA, Cat. #116-GS) and 1% penicillin/streptomycin (Thermo Fisher, Waltham, MA, USA, Cat. #15140122). All cells were grown in humidified incubator at 37°C with 5% CO_2_. 293FT cells were cultured in D-MEM high glucose and supplemented with 10% FBS, 0.1mM MEM non-essential amino acids, 6mM L-glutamine, 1mM MEM sodium pyruvate, 1% penicillin/streptomycin, and 500µg/ml geneticin.

### Reagents

To inhibit NO production, 2.5mM of L-NAME (Nω-Nitro-L-arginine methyl ester hydrochloride) was added to cell cultures (Sigma, St. Louis, MO, USA, Cat. # N5751). L-arginine, an NO synthase agonist, was used at the concentration of 2.5mM and obtained from Sigma (St. Louis, MO, USA, Cat. # A6969). NO donor SNAP (S-Nitroso-N-acetyl-DL penicillamin) was used at the concentration of 10µM (Sigma, St. Louis, MO, USA, Cat. # N3398). Galunisertib (LY2157299), a TGFβ inhibitor, was used at the concentration of 10µM and obtained from Cayman Chemical (Ann Arbor, MI, USA, Cat. # 15312).

### Antibodies

The following antibodies were used for western blotting and ICC/IF: anti-β actin (Sigma, A1978); anti-TGF Beta 1 (Abcam, ab92486); anti-LTBP1 (Abcam, ab78294); anti-phospho-Smad3 (Novus Biologicals, NBP1-77836); anti-phospho-TGFβR1 (Abcam, ab112095); anti-αSMA (Abcam, ab7817); anti-Vimentin (Thermo Fisher, PA5-86264); anti-Histone H3 (Cell Signaling Technologies, 4499S); anti-Smad 3 (Cell Signaling Technologies, 9523S); and anti-TGFβR1 (Abcam, ab31013).

### Western Blotting

Cell lysates were prepared by scraping cell culture dish with plastic policeman after addition of lysis buffer (25mM Tris pH 7.4, 150mM NaCl, 1mM EDTA, 1% NP-40 and 5% glycerol with addition of protease and phosphatase inhibitors). The lysate was spun down at 13,000 x g for 15 minutes at 4°C and the protein lysate was transferred to a new tube. To determine protein concentration, a BCA assay was performed (Thermo Fisher, 23225) according to manufacturer’s protocol. Reduced samples (by addition of 2-mercaptoethanol) were heated at 95°C for 5 minutes then loaded onto a 4-12% tris-glycine gradient gel (Thermo Fisher, XP04122BOX) and ran at 125 volts for 1-1.5 hours. Proteins were transferred to a PVDF membrane, and the PVDF membrane was briefly washed in TBS-T and blocked in 5% non-fat dry milk (NFDM) for 1 hour. The membrane was washed 3x for 5 minutes each and the primary antibody was added and incubated overnight at 4°C. The membrane was probed with HRP conjugated secondary antibody in 5% NFDM for 1 hour and washed again. The membrane was incubated with Super Signal West Dura substrate (Thermo Fisher, 34076) for 5 minutes and was imaged on a Syngene GBOX (model # Chemi XX6).

### Co-immunoprecipitation

Protein lysates were normalized between samples and pre-cleared with protein A/G agarose beads. Beads were removed, and the lysates were incubated with primary antibody overnight at 4°C. Protein A/G beads were blocked in 0.1% BSA and added to the lysate and incubated on a rotator at 4°C for 3-4 hours. The mixture was centrifuged and supernatant was removed. The beads were washed with lysis buffer and the sample was heated in SDS sample buffer to remove the protein. The samples were run on SDS-PAGE gel for visualization.

### Immunofluorescence staining and imaging

Cells were seeded and grown on 12mm round glass coverslips that were coated in PureCol bovine collagen (Advanced BioMatrix, 5005) according to the manufacture’s protocol. The coverslips were removed from the dish, added to a humidified chamber, and fixed with 4% paraformaldehyde for 15 minutes. The coverslip was washed in PBS-T three times (five minutes each) and permeabilized for 15 minutes with 0.5% triton-X100 and washed again. Blocking was performed by addition of 3% BSA for 1 hour followed by washing. Primary antibody in 3% BSA/ 0.1% saponin was added dropwise to the coverslips and incubated overnight at 4°C. The coverslips were washed 6x before addition of secondary antibody and left for 2 hours to incubate at room temperature. After washing, nuclei were stained with DAPI (0.5 ng/ml) and incubated for 10 minutes before a final wash. A drop of Fluoromount-G (Thermo Fisher, 00-4958-02) was added to each coverslip and mounted on a glass slide. Fluorescence imaging was performed on an Olympus IX70 microscope utilizing CellSens software.

### Prediction of sites of S-nitrosylation

The amino acid sequence of TGFβ mRNA CDS (NCBI Reference Sequence: NM_000660.7) was analyzed by GPS-SNO 1.0 software to predict which sites of the protein could undergo S-nitrosylation. Cysteine positions 355, 356 and 389 came back as positive targets that could harbor this modification.

### Site-directed mutagenesis of TGFβ

The QuikChange II Site-Directed Mutagenesis Kit (Agilent, 200524) was used according to manufacturer’s protocol. Primers were designed using Agilent’s QuikChange primer design online software and purchased from Thermo Fisher. The TGF beta 1 (NM_000660) Human Untagged Clone (Origene, SC119746) was used to perform site-directed mutagenesis on residues C355, C356 and C389. Each primer was designed to replace each three cysteine residues with an alanine residue. A PCR reaction was performed to create the new sequence with specific mutation sites. The PCR product was visualized on a 1% agarose gel to confirm the proper size. The product was transformed into XL-1 blue supercompotent cells to repair nicks in the DNA strand. The newly transformed XL-1 blue cells were spread onto agar petri plates containing 100µg/ml ampicillin and incubated upside down in a 37°C bacterial incubator overnight. Several colonies per plate were picked the next day and added to 5ml of LB broth containing ampicillin and grown again overnight at 37 °C while rocking at 220 RPM. The broth was collected and centrifuged at 5,000 x g to pellet the e-coli cells.

### Ligation into mammalian expression vector and transformation

A mini-prep (Qiagen, 27104) was conducted on pelleted cells from the transformation completed above following the mini-prep kit protocol. Purified plasmids were analyzed by DNA sequencing and confirmed the amino acid substitution for each site. To this DNA, PCR was performed to create specific restriction sites at the 5’ and 3’ ends of our gene of interest. The restriction sites added were for EcoRI and XbaI. A double restriction digest was performed to the newly synthesized sites to prepare the insert for ligation into the expression plasmid. After double digest was complete, the products were ran on a 1% agarose gel and the band of interest excised with a razor blade. The DNA was recovered by gel extraction utilizing the QIAquick Gel Extraction Kit (Qiagen, #28704). The DNA concentration of the insert was taken. In order to insert our gene of interest into a mammalian expression vector, we used the PCDF1-MCS2-EF1-Puro expression vector (System Biosciences, CD110B-1). We performed a restriction digest and gel extraction as above to the plasmid. We performed ligation of our insert and expression vector utilizing the T4 DNA ligase (New England Biolabs, M0202S) according to manufactures protocol. The ligation product was transformed into XL-1 blue supercompotent cells and plated as before. The following morning, we picked several colonies from each clone and grew them overnight in LB broth containing the appropriate antibiotics. We conducted another mini-prep and performed a double digest for EcoRI and XbaI to confirm the size of our insert.

### Transfection, lentivirus production, and transduction

For creation of pseudoviral particles containing mutant TGFβ1, 293FT cells were cultured and maintained according to the manufactures protocol. One day prior to transfection, 293FT cells were seeded to achieve 80% confluency the following day. The transfection was completed following purefection transfection reagent protocol (System Biosciences, LV750A). Briefly, reduced serum media (Thermo Fisher, #31985-062) was added to a tube along with purefection reagent and pPackF1 lentiviral packaging kit plasmids (System Biosciences, # LV100A-1) and either PCF1 (empty vector control), W/T, or point-mutated TGFβ1 plasmids. The mixture was incubated for 15 minutes and added dropwise to plates containing 293FT cells. The media containing pseudoviral particles was collected at 48 and 72 hours and syringe filtered to remove any cell debris. The viruses from each time point were pooled and concentrated using PEG-it virus precipitation solution (System Biosciences, LV810A) according to manufacture’s protocol. For transduction of target cells, MCF10A cells were seeded to achieve 50% to 70% confluency and the following day the virus particles were added to the dish along with TransDux virus transduction reagent (System Biosciences, LV850A). The media was changed after 4 days and antibiotic selection was performed using a puromycin concentration of 0.85µg/mL. The pSIH1-H1-siLuc-copGFP plasmid (System Bioscience, LV601PB) was used as a positive transduction control.

### S-nitrosylation assay

In order to determine the level of s-nitrosylated TGFβ, we used the Pierce S-Nitrosylation Western Blot Kit (Thermo Fisher, 90105) following the provided protocol. Briefly, MCF10A mutant and W/T TGFβ1 expressing cells were grown and harvested in HENS lysis buffer supplied with the kit. Protein concentrations were normalized between samples before beginning the assay. To each sample, methyl methanethiosulfonate (MMTS) was added to block free cysteine thiols. Protein precipitation was performed with ice-cold acetone to remove excess MMTS. The samples were centrifuged, acetone was removed, and the pellet was resuspended in HENS buffer. Sodium ascorbate and the labeling tag (iodo-TMT) were added and the samples were incubated. Protein precipitation was again performed and the pellet was resuspended in HENS buffer. In order to separate labeled from non-labeled proteins, samples were incubated with immobilized Anti-TMT Resin (Thermo Fisher, 90076). The eluted samples were run on SDS-PAGE for analysis.

### TGFβ measurement

To determine the amount of secreted TGFβ from the MCF10A cell line, we used the Human TGF beta 1 ELISA Kit (Abcam, ab108912). At day 1, cells were seeded in normal media. At day 2, roughly 24 hours later, the serum media was aspirated from the culture plates and serum-free media was added. The cells were left to grow until they reached approximately 90% confluency. The media was collected with a 10mL syringe and passed through a 0.45-µM filter to remove any cell debris. The media was concentrated by adding the sample to a 3kd cutoff protein concentrator and centrifuged. Once the media had decreased to roughly 1/10 its original volume, the fraction containing the concentrated protein was collected. Protein concentration was taken and samples were normalized against each other. ELISA was performed according to manufacturer’s protocol.

### Co-culture and conditioned media experiments

Briefly, fibroblasts cells were added on top of purecol coated coverslips in the bottom of a 12-well dish with a 1:1 ratio of MCF10A cell culture media to fibroblasts media. We placed the insert on top and added MCF10A cells along with more growth media. To treat the cells, drugs were added to the top insert. Coverslips were removed after treatment for 48 hours and cells were stained following the staining protocol. For conditioned media experiments, we drug-treated MCF10A cells and let them grow for an additional 24 hours. After 24 hours of incubation, media was collected from MCF10A culture plates with a syringe and filtered using a 0.45µM filter to remove cell debris. At the same time, we aspirated media from fibroblast culture plates and added the drug-treated MCF10A conditioned media to the plates. We repeated this once more for a total of two treatments. After conditioned media treatments on fibroblasts was complete, we harvested the coverslips and stained the cells following our protocol.

### RT-PCR

To determine mRNA levels of TGFβ in our mutant cell lines, we cultured cells and collected RNA utilizing an RNeasy Kit (Qiagen #74004) and created cDNA using SuperScript IV Reverse Transcriptase (ThermoFisher, #18090010) and followed the manufacturer’s protocol. We then performed PCR on cDNA and ran the products on a 1% agarose gel to visualize intensity of the products. (For primers see Table S1).

### Prediction of 3D monomeric and dimeric structures of TGFβ1

C-terminal 50 AA sequence of mature TGFβ1 (wild-type and different mutants) was input into PEP-FOLD3 software (34) which was freely available online. The resultant structures (PDB format) with the highest probability scores were further assembled into homodimers using SymmDoc software, which was also freely available online and could measure dimerization scores (35). These predicted 3D structures were visualized using UCSF Chimera software (50).

### Statistical methods

All the experiments were performed in replicates (n ≥ 3) and, unless otherwise indicated, two-tailed t-tests were performed to obtain the statistical significance of the mean difference. Statistical significance of the mean difference was tested by parametric tests using Graphpad Prism version 9. P-value of 0.05 or lower was considered significant and the average value is presented as the arithmetic mean ± SEM.

### Data, materials, and software availability

All study data are included in the article and/or SI Appendix.

## Acknowledgement

We thank Dr. Satyabrata Sinha in Furuta laboratory at Case Western Reserve University/MetroHealth Medical Center for manuscript review; and Drs. Gang, Ren, Xunzheng Zheng, and Vandana Sharma at University of Toledo Health Science Campus for technical help and constructive suggestions.

## Funding

This work was supported by the startup fund from University of Toledo Health Science Campus, College of Medicine and Life Sciences, Department of Cancer Biology to S.F; Ohio Cancer Research Grant (Project #: 5017) to S.F; Medical Research Society (Toledo Foundation, #206298) Award to S.F; American Cancer Society Research Scholar Grant (RSG-18-238-01-CSM) to S.F; and National Cancer Institute Research Grant (R01CA248304) to SF.

## Author contributions

Conceptualization, S.F. and J.L..; Methodology, S.F., and J.L; Formal Analysis, Investigation, J.L.; Data Curation, J.L.; Writing – Original Draft Preparation, J.L. and S.F.; Visualization, J.L.; Supervision, S.F.; Project Administration, S.F.; Funding Acquisition, S.F.

## Competing interests

The authors declare no competing interest.

## Supporting information

Appendix 01 (PDF)

## Appendix to

### Supplementary Table

**Table S1:**
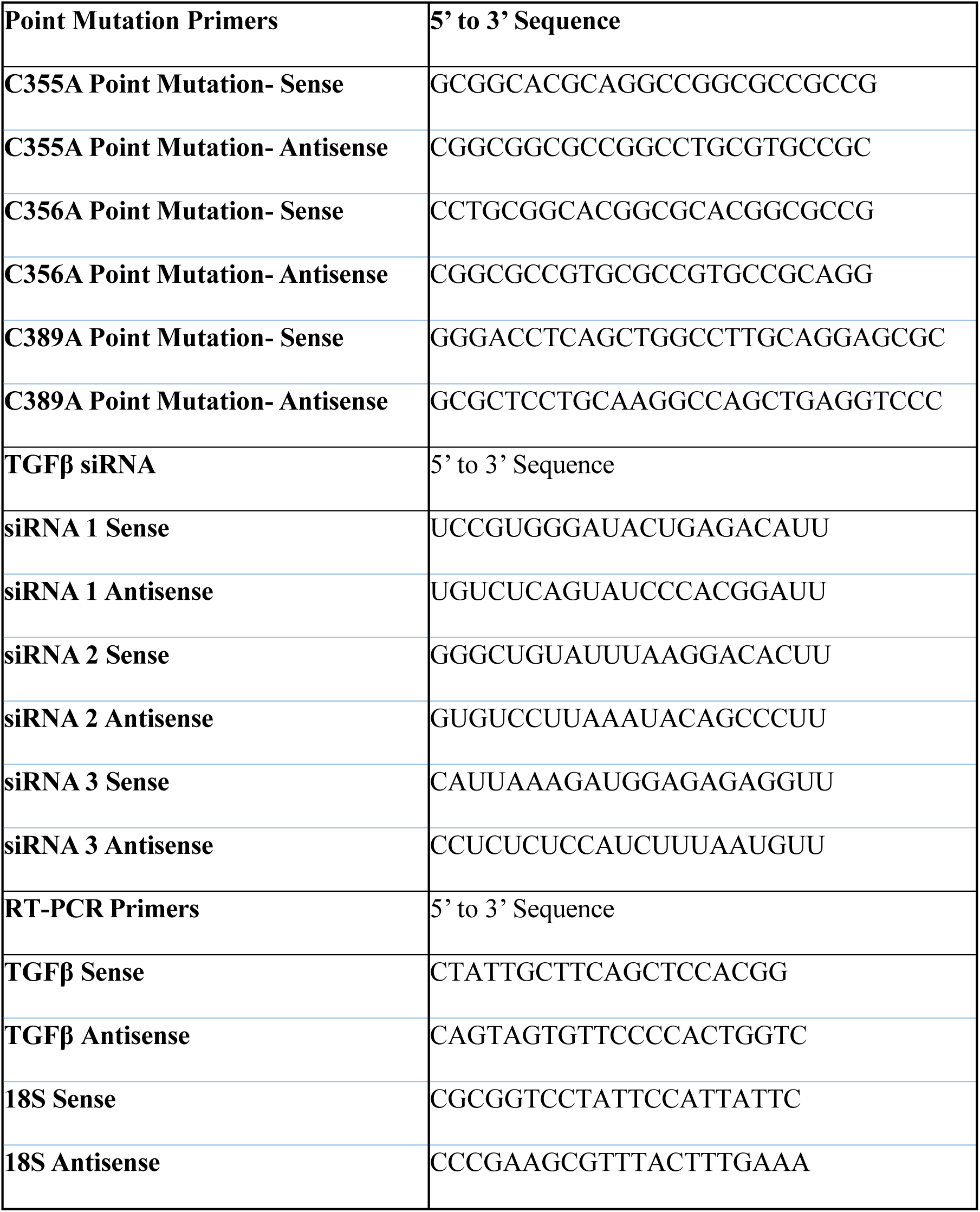
Primers and siRNAs used in experiments (See Materials and Methods).

### Supplementary Figure

**Figure S1:**
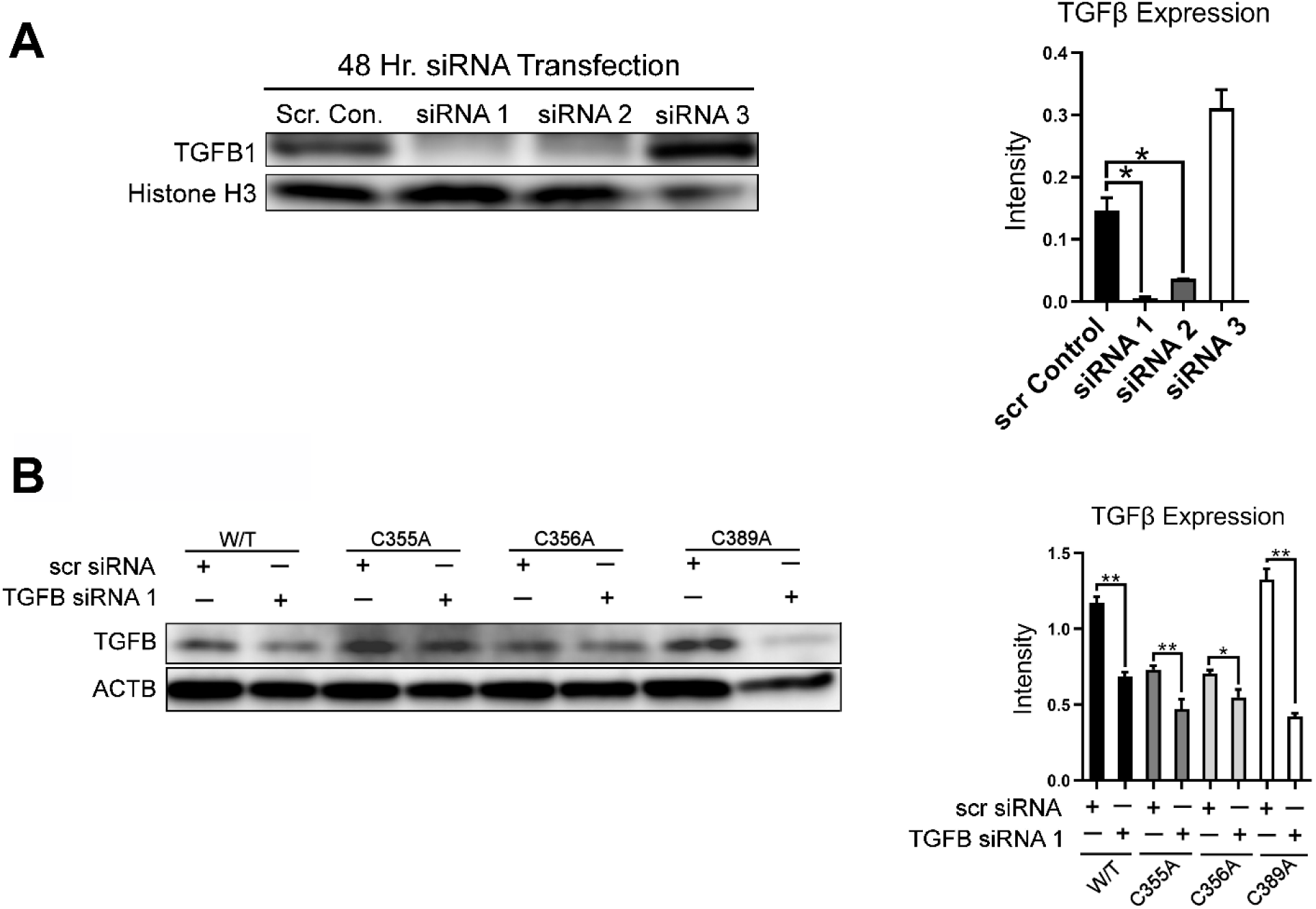
Treatment with siRNAs specifically removing the endogenous TGFβ1 (binding to either 3’ or 5’ UTR region of mRNAs) shows the levels of ectopic proteins. **A)** (Left) Western blot for the expression of TGFβ1 in MCF10A cells after treatment with three different siRNAs that target 3’ or 5’ UTR of the TGFβ1 transcripts. ACTB was used as an internal loading control. (Right) The intensities of TGFβ1 normalized against ACTB. Note that siRNA1 shows the strongest inhibitory effect. **B)** (Left) MCF10A cell lines expressing wild-type or SNO-defective mutants (C355A, C356A, or C389A) were treated with either scramble (scr) siRNA or siRNA1 (the most potent siRNA), and TGFβ1 levels were determined by Western blot. ACTB was used as an internal loading control. (Right) The intensities of TGFβ1 normalized against ACTB. Note that even after siRNA treatment, the majority of TGFβ1 proteins remained intact, indicating that they were ectopically expressed proteins. Error bars: ± SEM. *, and p ≤ 0.05. p >0.05 was considered significant.

## Notes

### Competing Interest Statement

The authors have declared no competing interest.

### Summary of Updates

Several typos have been corrected.

## References

1. Shao, Z.-M., Nguyen, M. & Barsky, S. H. (2000) Oncogene 19, 4337–4345.

2. Boyd, N. F., Martin, L. J., Yaffe, M. J. & Minkin, S. (2011) Breast Cancer Res 13, 223.

3. Foster, D. S., Jones, R. E., Ransom, R. C., Longaker, M. T. & Norton, J. A. (2018) JCI insight 3, e99911.

4. Sharma, V., Letson, J. & Furuta, S. (2022) Sci Signal 15, eabg3449.

5. Park, H. Y., Kim, J. H. & Park, C. K. (2013) Am J Pathol 182, 2147–54.

6. Moses, H. L., Roberts, A. B. & Derynck, R. (2016) Cold Spring Harb Perspect Biol 8.

7. Derynck, R., Akhurst, R. J. & Balmain, A. (2001) Nat Genet 29, 117–29.

8. Cole, K., Tabernero, M. & Anderson, K. S. (2010) Cancer Biomark 9, 177–92.

9. Silva, L. D., Parry, S., Reid, L., Keith, P., Waddell, N., Kossai, M., Clarke, C., Lakhani, S. R. & Simpson, P. T. (2008) The American Journal of Surgical Pathology 32.

10. Li, M., Luan, F., Zhao, Y., Hao, H., Zhou, Y., Han, W. & Fu, X. (2016) Experimental biology and medicine (Maywood, N.J.) 241, 1–13.

11. Ren, G., Zheng, X., Bommarito, M., Metzger, S., Walia, Y., Letson, J., Schroering, A., Kalinoski, A., Weaver, D., Figy, C., Yeung, K. & Furuta, S. (2019) Sci Rep 9, 6688.

12. Ren, G., Zheng, X., Sharma, V., Letson, J., Nestor-Kalinoski, A. L. & Furuta, S. (2021) Cancers (Basel) 13.

13. Zhao, Y., Tan, Y. S., Aupperlee, M. D., Langoh, r. I. M., Kirk, E. L., Troester, M. A., Schwartz, R. C. & Haslam, S. Z. (2013) Breast Cancer Res. 15, R100.

14. Wu Y, Zhang D & S, K. (2013) Breast Cancer Res Treat 137, 869–82.

15. Novosyadlyy, R., Lann, D. E., Vijayakumar, A., Rowzee, A., Lazzarino, D. A., Fierz, Y., Carboni, J. M., Gottardis, M. M., Pennisi, P. A., Molinolo, A. A., Kurshan, N., Mejia, W., Santopietro, S., Yakar, S., Wood, T. L. & LeRoith, D. (2010) Cancer Res. 70, 741–51.

16. Stoll, B. (2000) Int J Obes Relat Metab Disord 24, 527–33.

17. Stoll, B. (1999) Eur J Cancer 35, 1653–8.

18. Catsburg, C., Miller, A. B. & Rohan, T. E. (2015) Int J Cancer 136, 2204–9.

19. Han, H., Guo, W., Shi, W., Yu, Y., Zhang, Y., Ye, X. & He, J. (2017) Sci Rep. 7, 44877.

20. Martins, M. A., Catta-Preta, M., Mandarim-de-Lacerda, C. A., Aguila, M. B., Brunini, T. C. & Mendes-Ribeiro, A. C. (2010) Arch Biochem Biophys 499, 56–61.

21. Afsharm, M., Poole, J., Cao, G., Durazo, R., Cooper, R., Kovacs, E. J. & Sisson, J. H. (2016) Chest 150, 196–209.

22. Nyberg, M., Blackwell, J., Damsgaard, R., Jones, A. M., Hellsten, Y. & Mortensen, S. P. (2012) J Physiol 590, 5361–70.

23. Tessari, P., Cecchet, D., Cosma, A., Vettore, M., Coracina, A., Millioni, R., Iori, E., Puricelli, L., Avogaro, A. & Vedovato, M. (2010) Diabetes 59, 2152–9.

24. Williams, I., Wheatcroft, S. B., Shah, A. M. & Kearney, M. T. (2002) Int J Obes Relat Metab Disord 26, 754–64.

25. Bodis, S. & Haregewoin, A. (1994) Ann Oncol 5, 371–2.

26. Panza, J., A,, Casino, P. R., Kilcoyne, C. M. & Quyyumi, A. A. (1993) Circulation 87, 1468–74.

27. Fernando, V., Zheng, X., Walia, Y., Sharma, V., Letson, J. & Furuta, S. (2019) Antioxidants (Basel) 8.

28. Sharma, V., Fernando, V., Letson, J., Walia, Y., Zheng, X., Fackelman, D. & Furuta, S. (2021) Int J Mol Sci 22.

29. Imbalzano, K. M., Tatarkova, I., Imbalzano, A. N. & Nickerson, J. A. (2009) Cancer Cell International 9, 7.

30. Furuta, S. (2017) Trends in Cancer 3, 744–748 PMID: 29120749

31. Xue, Y., Liu, Z., Gao, X., Jin, C., Wen, L., Yao, X. & Ren, J. (2010) PLoS One 5, e11290.

32. Qu, Z., Meng, F., Bomgarden, R. D., Viner, R. I., Li, J., Rogers, J. C., Cheng, J., Greenlief, C. M., Cui, J., Lubahn, D. B., Sun, G. Y. & Gu, Z. (2014) J Proteome Res 13, 3200-11.

33. Huang, T., Schor, S. L. & Hinck, A. P. (2014) Biochemistry 53, 5737–49.

34. Lamiable, A., Thévenet, P., Rey, J., Vavrusa, M., Derreumaux, P. & Tufféry, P. (2016) Nucleic Acids Res 44, W449–54.

35. Schneidman-Duhovny, D., Inbar, Y., Nussinov, R. & Wolfson, H. J. (2005) Nucleic Acids Res 33, W363–7.

36. Duhovny D, N. R., Wolfson HJ (2002) Efficient Unbound Docking of Rigid Molecules. (Springer Verlag.

37. Zhang, C., Vasmatzis, G., Cornette, J. L. & DeLisi, C. (1997) J Mol Biol 267, 707–26.

38. Palmer, Z. J., Duncan, R. R., Johnson, J. R., Lian, L. Y., Mello, L. V., Booth, D., Barclay, J. W., Graham, M. E., Burgoyne, R. D., Prior, I. A. & Morgan, A. (2008) Biochem J 413, 479–91.

39. Wang, P., Liu, G. H., Wu, K., Qu, J., Huang, B., Zhang, X., Zhou, X., Gerace, L. & Chen, C. (2009) J Cell Sci 122, 3772–9.

40. Brunner, A. M., Marquardt, H., Malacko, A. R., Lioubin, M. N. & Purchio, A. F. (1989) Journal of Biological Chemistry 264, 13660–13664.

41. Liu, S., Ren, J. & ten Dijke, P. (2021) Signal Transduction and Targeted Therapy 6, 8.

42. López-Díaz, F. J., Gascard, P., Balakrishnan, S. K., Zhao, J., Del Rincon, S. V., Spruck, C., Tlsty, T. D. & Emerson, B. M. (2013) Mol Cell 50, 552–64.

43. Furler, R. L., Nixon, D. F., Brantner, C. A., Popratiloff, A. & Uittenbogaart, C. H. (2018) Cancers (Basel) 10.

44. Kolluru, G. K., Bir, S. C. & Kevil, C. G. (2012) Int J Vasc Med 2012, 918267.

45. Leopold, J. A. (2013) Hypertension 62, 1003–4.

46. Cox, T. R. & Erler, J. T. (2011) Dis Model Mech 4, 165–78.

47. Steppan, J., Sikka, G., Jandu, S., Barodka, V., Halushka, M. K., Flavahan, N. A., Belkin, A. M., Nyhan, D., Butlin, M., Avolio, A., Berkowitz, D. E. & Santhanam, L. (2014) J Am Heart Assoc 3, e000599.

48. Tanaka, H. (2019) Hypertension 74, 237–243.

49. Cohen, G., Steinberg, D. M., Keinan-Boker, L., Shaked, O., Goshen, A., Shimony, T., Shohat, T. & Gerber, Y. (2020) Mayo Clin Proc Innov Qual Outcomes 4, 115–125.

50. Pettersen, E. F., Goddard, T. D., Huang, C. C., Couch, G. S., Greenblatt, D. M., Meng, E. C. & Ferrin, T. E. (2004) J Comput Chem 25, 1605–12.

